# Leukemic mutation FLT3-ITD is retained in dendritic cells and disrupts their homeostasis leading to expanded Th17 frequency

**DOI:** 10.1101/2023.09.19.558512

**Authors:** Patrick A Flynn, Mark D Long, Yoko Kosaka, Jessica S Mulkey, Jesse L Coy, Anupriya Agarwal, Evan F Lind

**Affiliations:** Molecular Microbiology and Immunology, Oregon Health & Science University, Portland, OR 97201, USA; Department of Biostatistics and Bioinformatics, Roswell Park Comprehensive Cancer Center, Buffalo, NY 14263, USA; Cell, Developmental and Cancer Biology, Oregon Health & Science University, Portland, OR 97201, USA; Division of Oncological Sciences, Oregon Health & Science University, Portland, OR 97201, USA; Knight Cancer Institute, Oregon Health & Science University, Portland, OR 97201, USA

## Abstract

Dendritic cells (DC) are mediators of adaptive immune responses to pathogens and tumors. DC development is determined by signaling through the receptor tyrosine kinase Fms-like tyrosine kinase 3 (FLT3) in bone marrow myeloid progenitors. Recently the naming conventions for DC phenotypes have been updated to distinguish between “Conventional” DCs (cDCs) and plasmacytoid DCs (pDCs). Activating mutations of FLT3, including Internal Tandem Duplication (FLT3-ITD), are associated with poor prognosis for leukemia patients. To date, there is little information on the effects of FLT3-ITD in DC biology. We examined the cDC phenotype and frequency in bone marrow aspirates from patients with acute myeloid leukemia (AML) to understand the changes to cDCs associated with FLT3-ITD. When compared to healthy donor (HD) we found that a subset of FLT3-ITD+ AML patient samples have overrepresented populations of cDCs and disrupted phenotypes. Using a mouse model of FLT3-ITD+ AML, we found that cDCs were increased in percentage and number compared to control wild-type (WT) mice. Single cell RNA-seq identified FLT3-ITD+ cDCs as skewed towards a cDC2 T-bet**^-^**phenotype, previously shown to promote Th17 T cells. We assessed the phenotypes of CD4+ T cells in the AML mice and found significant enrichment of both Treg and Th17 CD4+ T cells. Furthermore, co-culture of AML mouse- derived DCs and naïve OT-II cells preferentially skewed T cells into a Th17 phenotype. Together, our data suggests that FLT3-ITD+ leukemia-associated cDCs polarize CD4+ T cells into Th17 subsets, a population that has been shown to be negatively associated with survival in solid tumor contexts. This illustrates the complex tumor microenvironment of AML and highlights the need for further investigation into the effects of FLT3-ITD mutations on DC phenotypes.

## Introduction

Dendritic cells (DCs) are cellular sentinels of the immune system that have evolved to detect pathogenic microorganisms, maintain tolerance to self, and prime T cells to their cognate antigens and contribute to protective immunity. In the decades after their formal description by Steinman and Cohn in 1979 [1, 2], the DC biology field has dramatically improved our understanding of both DC ontogeny and phenotype, which now includes cDC1, cDC2-subsets, and pDCs [3–8]. Among these findings is the key developmental signal for cDCs during hematopoiesis through the engagement of fms-like tyrosine kinase 3 (FLT3) by its ligand FLT3L and downstream activation of STAT3 [9]. Exogenous injection of FLT3L increases abundance of cDCs *in vivo* but there have not been reports of pathogenicity associated with high amounts of FLT3L [9–12].

Acute myeloid leukemia (AML) is a heterogeneous cancer of hematopoiesis that develops from the combination of two or more classes of mutations that affect proliferation, differentiation, and epigenetics of myeloid precursors in the bone marrow [13]. Activating mutations of FLT3 have a strong association with leukemias including AML [13–15]. It is the most common mutation found in AML patients, up to 30%, acquired by internal tandem duplications (FLT3-ITD) of the cytoplasmic domain of FLT3 [14, 16–20] and causes ligand independent signaling [16, 21]. There are few reports characterizing blood DC-subsets in both human leukemia and myelodysplastic syndromes [22, 23] and sample sizes are limited. Because of the genetic variability of the disease, it has been difficult to distinguish *bona fide* DCs from tumor cells with aberrant expression of DC-related proteins.

Mouse models to study AML have diverse mutation targets and cover both inducible and constitutive expression models with varying degrees of disease induction [24]. Despite the variety of models available, there is a paucity of DC- specific reports. It was shown in a non-leukemia model that the FLT3-ITD mutation produces functional DCs without pathogenic side effects but does result in increased DC abundance and moderate effects on CD4+ T cell phenotype [25].

To provide insight into the phenotypic and molecular changes in DCs during AML we investigated samples isolated from patients with AML and a genetically-engineered mouse model (GEMM) of AML that spontaneously develops disease. We first identified a disruption in the frequency of cDCs in the bone marrow of patients with AML segregated by their FLT3-ITD status, suggesting that some AML patients have increased output of DCs compared to healthy donors. We then utilized our GEMM AML mice and confirmed that the FLT3-ITD mutation causes significant expansion of cDCs both in the bone marrow and spleens of AML mice compared to FLT3-ITD**^-^** controls and healthy mice. Based on that finding we used single cell RNA-sequencing (scRNA-seq) to interrogate changes to AML cDCs at the transcript level and correlated the expression of surface proteins using antibody-derived tags (ADTs). We were able to identify *bona fide* cDCs transcriptionally and found an enrichment of the recently described T-bet**^-^** cDC2 that are efficient at polarizing naïve CD4+ T cells into Th17 cells. In the mice with AML, we observed elevated levels of blood serum cytokines that are permissive to Th17 polarization of CD4+ T cells. There is a significant enrichment of both Treg and Th17 phenotypes in the blood of AML mice, suggesting that AML DCs may be driving their polarization. To measure the impact of DCs on T cell skewing we used adoptive transfer studies to compare AML and WT healthy mice. We did not find Treg or Th17 polarization; however, AML hosts expanded and retained significantly more transferred OT-II cells compared to WT, suggesting that the increased DC frequency supports naïve OT-II T cell survival. Finally, we show that co-culturing naïve OT-II cells with AML DCs results in increased IL-17A production, suggesting that AML DCs play a role in polarizing Th17 cells in the presence of cognate antigen. Together, we propose that DC-progenitors retain the FLT3-ITD mutation which drives their significant expansion and leads to the activation of naïve CD4+ T cells that polarize into Th17 and Treg phenotypes. These T cell subsets would likely result in negative impacts to anti-tumor responses in AML.

## Materials and Methods

### Data availability statement

All single cell CITE-Seq data is accessible on Geo Datasets under the name “FLT3-ITD mutation expands dendritic cells and alters CD4+ T cells in acute myeloid leukemia” GSE238156. All other data available upon request.

### Ethics statement

Studies involving patients was collected under OHSU IRB 4422 Marc Loriaux, PI. in accordance with all local and institutional requirements. Animal studies were approved and conducted under OHSU IACUC protocol “Immune-based therapeutic approaches for acute myeloid leukemia” IP00000907 Evan Lind, PI.

### Human samples

Bone marrow aspirates and peripheral blood samples were separated by Ficoll density gradient centrifugation. All experiments were performed using liquid nitrogen stored samples. Antibody clones and source are listed in Table 1. Viability was determined by Zombie Aqua staining and doublets were gated out of analysis by FSC-A vs. FSC-H. Flow cytometry data were acquired on a BD LSRFortessa or Cytek Aurora and analyzed using FlowJo v10 software. All human sample experiments are approved under IRB protocol #00004422, “Pathogenesis of Acute Leukemia, Lymphoproliferative Disorder and Myeloproliferative Disorders” (PI: Marc Loriaux, MD, PhD). Informed consent was obtained from all patients.

### AML Murine Model

Mice expressing FLT3-ITD under the endogenous FLT3 promoter (strain B6.129-Flt3tm1Dgg/J, The Jackson Laboratory, stock no. 011112) [26] were crossed to mice with the TET2 gene flanked by LoxP sites (strain B6;129STet2tm1.1Iaai/J, The Jackson Laboratory, stock no. 017573) [27]. The Flt3ITD/Tet2 flox mice were then crossed to mice expressing Cre recombinase under the Lysm promoter (strain B6.129P2-Lyz2tm1(cre)Ifo/J, The Jackson Laboratory, stock no. 004781). The Flt3ITD/Tet2/LysMCre mice were bred to mice with the TP53 gene flanked by LoxP sites (strain B6.129P2-Trp53tm1Brn/J, The Jackson Laboratory, stock no. 008462). All breeding animals were purchased from The Jackson Laboratory. All mice used in these experiments were bred as heterozygous for FLT3-ITD and LysCre but homozygous for TET2 and TP53. All mouse experiments were performed in accordance with the OHSU Institutional Animal Care and Use Committee protocol IP00000907. No inclusion or exclusion criteria were used on the animals with correct genotype. Mice were selected and assigned to groups randomly while maintaining a 50% male 50% female ratio per experiment. No blinding was performed. Average age of mice used for in vivo studies was 30 weeks.

### *Ex vivo* cell preparation

Spleens were harvested from mice and mechanically dissociated using frosted microscope slides then rinsed with 1x PBS. Single cell suspensions were passed through 70-µm cell strainers and red blood cells were then lysed with ammonium chloride-potassium (ACK) lysis buffer. Cells were counted with hemacytometer and 3–5 × 10^6^ cells were used per antibody-staining reaction. For experiments requiring enrichment of splenic cDCs, cells were enriched from total spleen cells with MojoSort™ Mouse CD11c Nanobeads (BioLegend Cat: 480078) according to manufacturer’s protocol. After positive selection cells were counted via hemacytometer to assess viability and cell count with purity assessed via flow cytometry. For adoptive transfer and co-culture experiments naïve OT-II cells were magnetically enriched with MojoSort™ Mouse CD4 Naïve T Cell Isolation Kit (BioLegend Cat: 480040) according to manufacturer’s protocol. After positive selection cells were counted via hemacytometer to assess viability and cell count with purity assessed via flow cytometry.

### Flow cytometry staining

Bone marrow, blood, or splenocytes were processed and subjected to red blood cell lysis by ACK before counting via hemacytometer. Cells were resuspended in PBS and stained at 4°C with 100 µL 1:500 Zombie Aqua viability dye (BioLegend, Cat# 423102) and 1:200 mouse FC block (TruStain FcX, BioLegend Cat# 101320) for 15 min, covered from light. Cells were pelleted for 5 minutes at 300xg. 100 µL of cell surface staining antibody cocktail was added directly on top of the cells and stained on ice for 30 min in FACS buffer (1X PBS, 1.5% calf serum, 0.02% sodium azide, 2mM EDTA). For intracellular staining, the cells were then washed with FACS buffer, permeabilized, and stained for intracellular targets according to the manufacturer’s protocol (eBioscience FOXP3 Transcription Factor Staining Buffer Set, Cat# 00-5523-00), then resuspended in FACS buffer before analyzing on either a BD Fortessa or Cytek Aurora flow cytometer. Data were analyzed using FlowJo software.

### Adoptive transfer experiments

Spleens from CD45.2^+^ OT-II mice [28] were harvested from mice and prepared for naïve OT-II cell transfer as described above before transfer to AML CD45.1^+^ or WT CD45.1^+^ recipient mice. On Day 0 200,000 naïve OT-II cells were intravenously injected into recipient mice. On day 1 soluble OVA protein (200 µg) was injected into recipient mice. On Day 11 spleens from recipient mice were harvested and splenocytes were prepared for flow cytometry staining to asses OT-II populations for cell number and frequency as described above.

### Antibodies

**Table 1.** Human antibody list.

| **<u>Antigen</u>** | **<u>Fluor</u>** | **<u>Clone</u>** | **<u>Catalogue</u>** | **<u>Vendor</u>** |
| --- | --- | --- | --- | --- |
| AXL | Alexa Fluor 350 | 108724 | FAB154U-100G | BD Bio |
| XCR1 | BV421 | S15046E | 372610 | BioLegend |
| CD11b | Pacific Blue | ICRF44 | 558123 | BD Bio |
| Zombie Aqua |  |  | 423102 | BioLegend |
| CD45 | BV605 | HI30 | 304042 | BioLegend |
| CD123 | BV650 | 6H6 | 306020 | BioLegend |
| CD33 | BV711 | WM53 | 303424 | BioLegend |
| CD19 | BV786 | HIB19 | 302240 | BioLegend |
| CD11c | Alexa Fluor 488 | Bu15 | 337236 | BioLegend |
| CD3e | PerCP | OKT3 | 317338 | BioLegend |
| CD34 | PerCP-Cy5.5 | 561 | 343612 | BioLegend |
| CLEC10A | PE | H037G3 | 354704 | BioLegend |
| CD56 | PE Dazzle 594 | HCD56 | 318348 | BioLegend |
| HLA-DR | PE-Cy5 | L243 | 307608 | BioLegend |
| CD1c | PE-Cy7 | L161 | 331516 | BioLegend |
| CD38 | Alexa Fluor 647 | HH2 | 303514 | BioLegend |
| CD14 | Alexa Fluor 700 | HCD14 | 325614 | BioLegend |

**Table 2.** Mouse antibody list.

| **Antigen** | **Fluor** | **Clone** | **Catalogue** | **Vendor** |
| --- | --- | --- | --- | --- |
| CD45 | BUV496 | 30-F11 | 749889 | BD Bio |
| CD86 | BUV737 | GL1 | 741737 | BD Bio |
| I/A-I/E | BV421 | M5/114.15.2 | 107631 | BioLegend |
| CD80 | BV650 | 16-10A1 | 104732 | BioLegend |
| CD317/BST2/PDCA-1 | BV711 | 927 | 104732 | BioLegend |
| XCR1 | BV785 | ZET | 148225 | BioLegend |
| CD103 | Alexa Fluor 488 | 2_E7 | 121408 | BioLegend |
| CD135 | PE | A2F10 | 135306 | BioLegend |
| CD11b | PE-Cy7 | M1/70 | 101216 | BioLegend |
| F4/80 | APC | BM8 | 123116 | BioLegend |
| CD25 | PE-Cy7 | PC61 | 102026 | BioLegend |
| CD45.1 | BV711 | A20 | 110739 | BioLegend |
| CD11c | Alexa Fluor 488 | N418 | 117311 | BioLegend |
| Ly6G | PerCP-Cy5.5 | 1A8 | 127616 | BioLegend |
| CD3e | PE | 145-2C11 | 100308 | BioLegend |
| SIRPα | PE-Dazzle 594 | P84 | 144016 | BioLegend |
| CD19 | PE-Dazzle 594 | 6D5 | 115554 | BioLegend |
| GR-1 | BV605 | RB6-8C5 | 108439 | BioLegend |
| CD3e | BV421 | 145-2C11 | 100336 | BioLegend |
| CD317/BST2/PDCA-1 | BV711 | 927 | 127039 | BioLegend |
| CD11b | APC | M1/70 | 101212 | BioLegend |
| Ly6C | Alexa Fluor 700 | HK1.4 | 128024 | BioLegend |
| CD62L | APC-Cy7 | MEL-14 | 104428 | BioLegend |
| CD86 | PE-Cy7 | PO3 | 105116 | BioLegend |
| CD86 | BV650 | GL-1 | 105035 | BioLegend |
| XCR1 | BV650 | ZET | 148220 | BioLegend |
| CD86 | APC-Cy7 | GL-1 | 105030 | BioLegend |
| B220 | PerCP-Cy5.5 | RA3-6B2 | 103236 | BioLegend |
| I/A-I/E | PE | M5/114.15.2 | 107608 | BioLegend |
| IFNγ | Pacific Blue | XMG1.2 | 505818 | BioLegend |
| IL-4 | BV605 | 11B11 | 504126 | BioLegend |
| I/A-I/E | BV650 | M5/114.15.2 | 107641 | BioLegend |
| IL-17A | BV711 | TCH11-18H10.1 | 506941 | BioLegend |
| FOXP3 | PE-CY7 | FJK.16S | 25-5773-82 | eBioscience |
| GATA3 | APC | 16E10A23 | 653805 | BioLegend |
| T-bet | BV786 | O4-46 | 564141 | BioLegend |
| RORγt | PE | B2D | 12-6981-82 | eBioscience |
| CD25 | APC-R700 | PC61 | 565134 | BD Bio |
| CD45.2 | APC-Cy7 | 104 | 109824 | BioLegend |
| CD11c | BV421 | N418 | 117330 | BioLegend |
| CD8a | Pacific Blue | 53-6.7 | 100725 | BioLegend |
| CD44 | BV605 | IM7 | 103407 | BioLegend |
| NK1.1 | BV650 | PK136 | 108736 | BioLegend |
| CD19 | BV711 | 6D5 | 115555 | BioLegend |
| CD3e | Alexa Fluor 488 | 145-2C11 | 100321 | BioLegend |
| I/A-I/E | PerCP | M5/114.15.2 | 107624 | BioLegend |
| CD4 | PerCP-Cy5.5 | RM4-4 | 116012 | BioLegend |
| Zombie Aqua |  |  | 423102 | BioLegend |

**Table 3.** ADT list.

| **Antigen** | **Clone** | **Catalogue** | **Vendor** |
| --- | --- | --- | --- |
| TotalSeq™-A0002 anti-mouse CD8a | 53-6.7 | 100773 | BioLegend |
| TotalSeq™-A0103 anti-mouse/human CD45R/B220 Antibody | RA3-6B2 | 103263 | BioLegend |
| TotalSeq™-A0106 anti-mouse CD11c Antibody | N418 | 117355 | BioLegend |
| TotalSeq™-A0014 anti-mouse/human CD11b Antibody | M1/70 | 101265 | BioLegend |
| TotalSeq™-A0117 anti-mouse I-A/I-E Antibody | M5/114.15.2 | 107653 | BioLegend |
| TotalSeq™-A0200 anti-mouse CD86 Antibody | GL-1 | 105047 | BioLegend |
| TotalSeq™-A0201 anti-mouse CD103 Antibody | 2E_7 | 121437 | BioLegend |
| TotalSeq™-A0212 anti-mouse CD24 Antibody | M1/69 | 101841 | BioLegend |
| TotalSeq™-A0422 anti-mouse CD172a (SIRPα) Antibody | P84 | 144033 | BioLegend |
| TotalSeq™-A0563 anti-mouse CX3CR1 Antibody | SA011F11 | 149041 | BioLegend |
| TotalSeq™-A0568 anti-mouse/rat XCR1 Antibody | ZET | 148227 | BioLegend |
| TotalSeq™-A0811 anti-mouse CD317 (BST2, PDCA-1) Antibody | 927 | 127027 | BioLegend |
| TotalSeq™-A0849 anti-mouse CD80 Antibody | 16-10A1 | 104745 | BioLegend |
| TotalSeq™-A0903 anti-mouse CD40 Antibody | 3/23 | 124633 | BioLegend |
| TotalSeq™-A1010 anti-mouse CD205 (DEC-205) Antibody | NLDC-145 | 138221 | BioLegend |
| TotalSeq™-A0093 anti-mouse CD19 Antibody Antibody | 6D5 | 115559 | BioLegend |
| TotalSeq™-A0981 anti-mouse TCR Vα2 Antibody | B20.1 | 127831 | BioLegend |
| TotalSeq™-A0013 anti-mouse Ly-6C Antibody | HK1.4 | 128047 | BioLegend |
| TotalSeq™-A0015 anti-mouse Ly-6G Antibody | 1A8 | 127655 | BioLegend |
| TotalSeq™-A0098 anti-mouse CD135 Antibody | A2F10 | 135316 | BioLegend |
| TotalSeq™-A0105 anti-mouse CD115 (CSF-1R) Antibody | AFS98 | 135533 | BioLegend |

### LegendPlex Blood Serum Assay

Peripheral blood from WT and AML was collected via retro-orbital sources. Blood was collected in Sarstedt Microvette® 500 Serum Gel tubes and centrifugated at 10,000 x G for 5 minutes at room temperature to remove cells from the serum. Aliquots of the remaining serum were stored at -80°C before assayed. On day of being assayed aliquots were thawed slowly on ice before being tested using the flow cytometry based LEGENDplex™ Mouse Inflammation Panel (BioLegend Cat: 740446) according to the manufacturer’s instructions for sample processing. Samples were measured on a BD Fortessa according to the manufacturer’s instructions for calibration and data collection. Data was analyzed using the online resource from BioLegend (https://legendplex.qognit.com).

### ADTs and Single cell RNA-seq

Spleens from WT and AML mice were harvested and processed into filtered single cell suspensions as described above. After counting, cells were enriched for DC populations using negative selection EasySep™ Mouse Pan-DC Enrichment Kit II (StemCell Cat: 19863) according to the manufacturer’s protocol. After magnetic enrichment unlabeled cells were counted and aliquoted into 1x10^6^ total cells for TotalSeq™-A (BioLegend, custom panel) ADT staining. Cells were incubated with 1:200 TruStain FcX™ PLUS (BioLegend Cat: 156604) in a final volume of 50 µL 1X FACS buffer for 10 minutes at 4°C. ADT 2X cocktail master mix was prepared for a final dilution of approximately 1 µg per ADT as follows: 1.8 µL per ADT for a total volume of 37.8 µL of just ADTs. 412.2 µL of 1X FACS buffer added to the ADTs for a final volume of 450 µL 2X ADT master mix cocktail. Add 50 µL of the master mix to each sample for a final volume of 100 µL. Incubate samples for 30 minutes at 4°C. Add 100 µL 1X FACS buffer and pellet cells for 5 minutes at 300xg. Wash two more times to remove any unbound ADTs. Resuspend cell pellets at 1,000 cells per µL with 1X PBS 0.04% BSA. Samples were transferred to the OHSU Massively Parallel Sequencing Shared Resource (MPSSR) for 10X Genomics Chromium CITE-seq library and scRNA-seq cDNA library preparation according to 10X Genomics protocols. Each sample was sequenced on its own lane on the chip with no multiplexing. Each sample was sequenced with a target number of 10,000 cells per sample at a reads of 20,000 per cell for an approximate read depth of 200 million reads.

### CITE-seq Analysis

Raw sequence data demultiplexing, barcode processing, alignment (mm10) and filtering for true cells were performed using the Cell Ranger Single-Cell Software Suite (v6.0.2), yielding 89,537 cells (WT: 11,326 cells per sample, AML: 11,059 cells per sample) with a mean of 23,788 reads/cell (91.38% mapping rate), median of 1,676 genes/cell, 20,037 total unique detectable genes, and 5,204 median UMI counts/cell. Subsequent filtering for high quality cells, and downstream analyses were performed using Seurat (v4) [29] (Supplementary Figure 3A-B). Genes expressed in less than 3 cells and cells that express less than 300 genes were excluded from further analyses. Additional filtering of cells was determined based on the overall distributions of total RNA counts (< 80,000) and the proportion of mitochondrial genes (< 10%) detected to eliminate potential doublets and dying cells, respectively. Additional detection of doublets was performed using Scrublet [30] using a cut-off threshold based on total distribution of doublet scores (doublet score < 0.2). Quantification of mitochondrial and ribosomal gene expression was calculated using the PercentageFeatureSet function, using gene sets compiled from the HUGO Gene Nomenclature Committee database. Cell cycle phase scoring was accomplished against normalized expression via the CellCycleScoring function using mouse genes orthologous to known cell cycle phase marker genes. Ultimately, 74,198 high quality cells (WT: 37,033 cells, AML: 37,165 cells) across samples were included in downstream analyses. Ambient RNA correction was performed using SoupX [31]. Normalization and variance stabilization of gene expression data were conducted using regularized negative binomial regression (sctransform) implemented with Seurat. Normalization of ADT abundances was accomplished via the centered log-ratio method. Principal component analysis (PCA) was performed on normalized data and optimal dimensionality of the dataset was decided by examination of the Elbow plot, as the total number of PCs where gain in cumulative variation explained was greater than 0.1% (PCs = 41). Unsupervised cluster determination was performed against a constructed SNN graph via the Louvain approach using a resolution of 0.08. UMAP was applied for non-linear dimensionality reduction to obtain a low dimensional representation visualization of cellular states. Differential expression between clusters or samples was determined using the MAST method [32] via the FindMarkers function, using a minimum expression proportion of 25% and a minimum log fold change of 0.25. Mean expression of markers found within each cluster or cell annotation were used for subsequent analyses including dotplot visualization. Major cell lineages were determined using SingleR [33] using gene sets derived from the ImmGen database. Reference based mapping of the DC compartment was performed against a previously published dataset characterizing DC heterogeneity in mouse spleen (Brown et. al. [5]). Raw counts were obtained from the Gene Expression Omnibus (GSE137710) and re-processed utilizing a Seurat based pipeline described above. Reference based mapping and label transfer of previously annotated cells was performed using the FindTransferAnchors and MapQuery functions implemented within Seurat. DC-specific annotations were further refined using curated marker assessment. Unbiased T-cell phenotype annotation was performed using ProjecTILs [34] and refined using knowledge-based assessment of canonical gene markers.

### Statistics

A 2-sample Student t test with unequal variances was used to test for a statistical difference between 2 groups or a 1-sample Student t test was used to test for a statistical difference in the percent change from 0% where indicated. Unless otherwise indicated, all hypothesis tests were 2-sided, and a significance level of 0.05 was used. Ordinary One-Way ANOVA used to test-for statistical differences between 3 groups. Statistics and Graphs were generated by using Prism 9 software (GraphPad Software https://www.graphpad.com/).

## Results

### Bone marrow samples from patients with AML exhibit disrupted development of cDCs

Bone marrow aspirates from patients with AML were analyzed by flow cytometry to measure frequencies of cDCs based on their expression of high levels of CD11c and HLA-DR on their surface (Figure 1A) after gating out non-DC immune cells (Supplemental 1A). We found that DCs from patients with AML showed a heterogeneous phenotype compared to healthy donors (HD). For a subset of patients, the frequency of cDCs as a proportion of CD45+ cells was increased compared to HD and this trend was even more pronounced in the samples from FLT3-ITD+ patients (Figure 1B). We then sought to investigate if there were any changes to the major cDC subtypes. We therefore assessed the expression of CD1c and XCR1 to identify cDC2 and cDC1, respectively (Figure 1C). The frequency of XCR1+ cDC1s was significantly lower in both AML patient groups compared to HD (Figure 1D). The frequency of CD1c+ cDC2 was significantly lower in FLT3-ITD+ patient samples but there was no difference between HD and FLT3-WT AML (Figure 1E). Unexpectedly the CD1c XCR1 double-negative population was significantly higher in the FLT3-ITD+ AML group compared to HD and FLT3-WT (Figure F). It has been recently appreciated that cDC2 can be further subtyped by their expression of cell surface CLEC10A as an alternative to intracellular T-bet staining [5] and therefore measured CLEC10A expression on CD1c+ cDC2s (Figure 1G). We observed that the frequency of CLEC10A+ CD1c+ cells was significantly lower in the FLT3-ITD+ AML group compared to HD and FLT3-WT AML (Figure 1H) and the same was true for the CLEC10A- cDC2s (Figure 1I). Our flow cytometry data is similar to what was found from bulk-RNA sequencing analysis of “DC-Like” signatures from the BEAT AML dataset (n=531), where subsets of AML patients with FLT3-ITD had increased “DC-Like” score compared to HD (Supplemental 1B) [14, 35]. Taken together, this data suggests that in some patients with FLT3-ITD+ AML, the homeostasis of bone marrow cDCs is disrupted and is characterized by a previously unreported expansion of double-negative XCR1-cDC1- poorly-differentiated cDCs.

**Figure 1.**
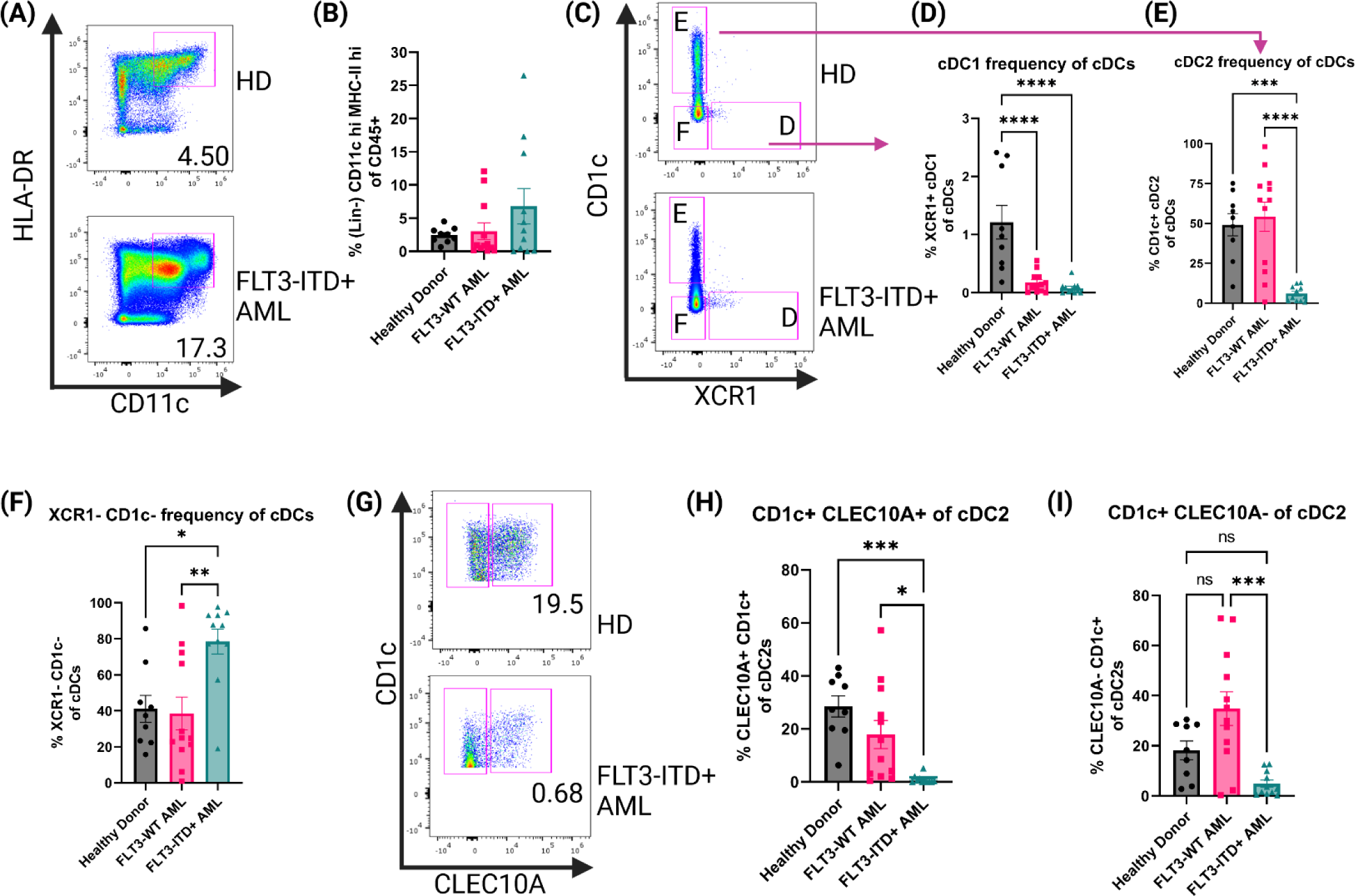
Human Bone Marrow Phenotyping Reveals Disruption of cDC Populations A. Representative dot plots showing human bone marrow cDCs. cDCs defined as Lin(CD3e, CD19, CD56, CD14, AXL, CD123)-CD45^+^CD11c^+^HLA-DR^+^. B. Summary graph for (A). Each symbol is one human sample. HD n=9, FLT3-WT AML n=12, FLT3-ITD^+^ AML n=11. C. Representative dot plots showing cDC1 and cDC2 subsets of the cDCs identified in (A). cDC1 defined as CD1c-XCR1^+^ and cDC2 defined as XCR1-CD1c^+^. (D-F) Summary graphs for (C). Each symbol is one human sample. HD n=9, FLT3-WT AML n=12, FLT3-ITD^+^ AML n=11. (G) Representative dot plots showing CLEC10A expression of cDC2s. (H-I) Summary dot plots of CLEC10a^+^ and CLEC10a- cDC2s as a frequency of total cDCs (A).

### Changes in cDCs in a FLT3-ITD+ mouse mode of AML

To better understand how the FLT3-ITD mutation in AML affects cDCs, we characterized cDCs in a mouse model of AML. Our lab has previously published work using AML mice that harbor the FLT3-ITD mutation and spontaneously develop AML to interrogate T cell dysfunction [36, 37]. Here we utilized mice that express one copy of FLT3-ITD under the endogenous FLT3 promoter, have a homozygous loss of TET2 and a homozygous loss of p53 using Cre-mediated deletion of LoxP-flanked alleles (AML mice) [21, 27, 38, 39]. This model is similar to others who reported combining FLT3-ITD with loss of TET2, but in our model the mutations are limited to the myeloid compartment by use of LysM-Cre, thereby retaining a wild-type lymphoid compartment [40]. This combination of mutations produces spontaneous AML, as evidenced by splenomegaly (Figure 2A&B) and increased CD11b+ cell frequency in both the spleen and the blood (Figure 2C&D). Furthermore, AML mice succumb to disease and die prematurely (Figure 2E). We then measured populations of DCs using the canonical surface markers CD11c and MHC-II (Figure 2F) which are expressed at high levels by cDCs compared to other myeloid subsets. We found that in both compartments, AML mice exhibited significantly increased cDCs (Figure 2G&H). When comparing the frequency of cDCs in the bone marrow and spleen to healthy WT controls we found that cDCs were significantly more abundant in AML mice (Figure 2I). The phenotype observed in AML mice was not observed in littermates that have wild-type FLT3 but still harbored loss of both TET2 and p53 (Supplemental Figure 2), supporting our hypothesis that FLT3-ITD would lead to increased abundance of cDCs *in vivo*. To confirm that FLT3-ITD has constitutive signaling in cDCs, we performed intracellular flow cytometry to measure phosphorylated FLT3 (pFLT3) using an antibody that recognizes Y591 of the intracellular domain of FLT3 (Figure 2J) [41, 42]. When comparing the geometric mean fluorescent intensity (gMFI) between WT and AML we found that circulating cDCs in AML blood have more pFLT3, as expected in mice with a FLT3-ITD mutation (Figure 2K). Taken together, our AML mouse model displays significant expansion of cDCs as a result of FLT3-ITD.

**Figure 2.**
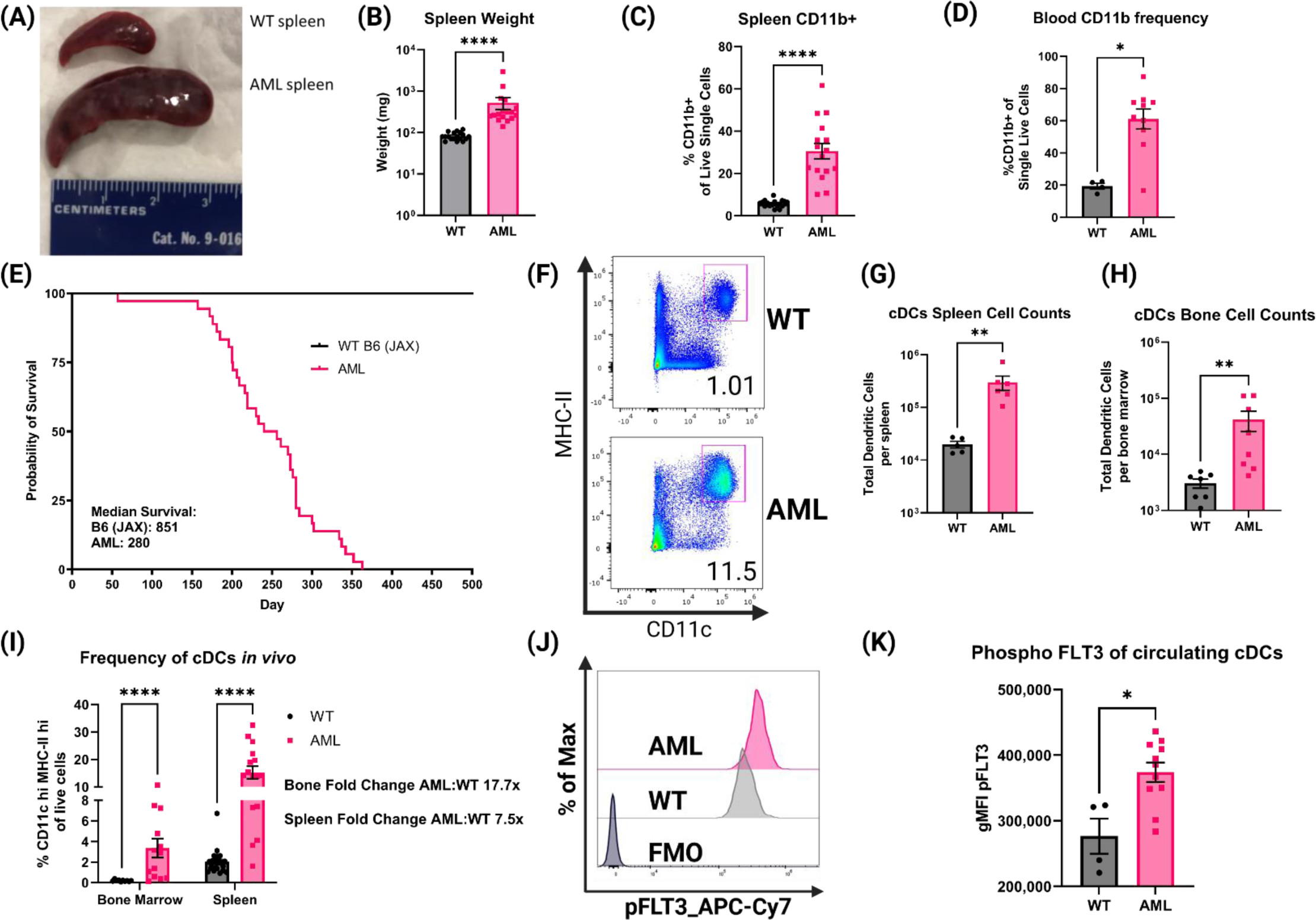
AML mice have increased frequencies of myeloid cells and cDCs. A. Comparison of dissected spleens from healthy WT mice and diseased AML mice. Ruler for scale (cm). B. Summary graph of spleen wet weight for WT and AML mice. Each symbol is one mouse. WT n=18 AML n=17 C. RBC lysed single-cell suspensions of WT and AML mice spleens were stained with monoclonal antibodies for known immune cell markers. Summary graph for frequency of CD19-CD3e-CD11b^+^ splenocytes. Each symbol is one mouse. WT n=17 AML n=16. D. RBC lysed single-cell suspensions of WT and AML PBMCs were stained with monoclonal antibodies for known immune cell markers. Summary graph for frequency of CD19-CD3e-CD11b^+^ PBMCs. Each symbol is one mouse. WT n=4 AML n=10 E. Survival curves of reference C57BL/6J mice from Jackson Laboratories (B6) and our mouse model AML mice. F. Representative dot plots showing mouse splenic cDCs. cDCs defined as Lin(Ly6C, Ly6G, F4/80, CD3e, CD19, NK1.1)- CD45^+^CD11c^hi^MHC-II^hi^. G. Summary graph of cDC cell counts per spleen of WT and AML mice. Each symbol is one mouse. WT n=5 AML n=6 H. Summary graph of cDC cell counts per bone marrow of WT and AML mice. Each symbol is one mouse. WT n=5 AML n=6 I. Summary graph of (F). Each symbol is one mouse. WT bone marrow n=10 WT spleens n=23. AML bone marrow n=13 AML spleens n=16. J. Representative histograms of phospho-FLT3 (pFLT3) staining in cDCs as measured by flow cytometry. RBC lysed single-cell suspensions of WT and AML PBMCs were stained with monoclonal antibodies for known immune cell markers at the surface and were stained for pFLT3 in the cytoplasm. K. Summary graph showing geometric mean fluorescent intensity (gMFI) pFLT3 staining of blood circulating cDCs. Each symbol is one mouse. WT n=4 AML n=11.

### Single-cell profiling of splenic cDCs in the context of FLT3-ITD AML

After confirming the phenotype of DCs in our AML mice, we interrogated how DCs are altered in the FLT3-ITD AML environment at the single-cell level using paired transcriptome and cell surface protein profiling (CITE-seq). In total, 74,198 high-quality cells were profiled across samples (WT: n = 4, AML: n = 4) after quality assessment and filtering procedures (Supplemental Figure 3A-B). Unsupervised cell clusters were annotated to major cell lineages using a combination of supervised and knowledge-based cell annotation procedures, and allowed for the identification of bona fide cDCs from other immune cell subsets as well as any possible AML-tumor related myeloid cells from spleen tissue (Supplemental Figure 3D-E). As expected, splenic cells captured from AML mice had a significantly higher proportion of DCs relative to WT mice (Supplemental Figure 4). Cell surface protein abundances measured by Antibody Derived Tags (ADTs) confirmed major cell lineages determined by transcriptome-based annotations (Supplemental Figure 4D), and strongly correlated with corresponding transcripts across cells (Supplemental Figure 5). Having isolated the DC compartment from other immune cells in the WT and AML splenic environment, we next sought to further define DC phenotype heterogeneity and how it is altered in AML. Recent single-cell profiling of DCs has established further heterogeneity than previously thought, for instance that cDC2 subsets can be segregated by their expression of T-bet [5]. To identify DC subtypes within our mouse splenic cells, we implemented a referenced based classification approach utilizing single-cell data derived from mouse splenocytes reported by Brown *et. al.* [5] (Supplemental Figure 6) to help in delineating DC phenotypes in our data. Using this approach paired with subsequent marker based assessment, we detected 9 distinct DC subsets including cDC1, T-bet- and Tbet+ cDC2, migratory *Ccr7* expressing DCs (CCR7+ DC), plasmacytoid DCs (pDC) and Siglec-H expressing pDC precursors (Siglec-H DC), monocyte-like DCs, as well as proliferative DC subsets which were strongly supported by canonical marker assessment (Figure 3A-C). Notably, upon examination of DC subtype proportions between WT and AML splenocytes, we observed a strong and significant shift in cDC2 subsets with AML splenocyte cDC2s being dominated by a Tbet- phenotype, while WT cDC2s maintained a Tbet+ phenotype (Figure 3D-E). The expression of *Clec10a* and *Cd209a* were highly expressed on T-bet- cDC2s (Figure 3B) in accordance with data published [5]. Surface protein detection by ADT confirmed that T-bet- cDC2s had low expression of CD80 but high expression of CD172a and CD11b as expected (Figure 3C). Amongst other DC subtypes, there was no observed difference in the proportion of cDC1s between AML and WT, but significant decreases in pDCs and CCR7+ DCs, and increase in monocyte-like DCs were also observed in AML splenocytes, although to a lesser extent than the shift in Tbet-/+ cDC2s. T-bet- cDC2s were previously reported to preferentially skew naïve CD4+ T cells into Th17s *in vitro* [5]. Th17 cells have been shown to be elevated in AML patients and have a complicated role in cancer [43, 44]. In summary, our data shows that *bona fide* DCs are present in AML and they are skewed towards T-bet- cDC2 phenotype.

**Figure 3.**
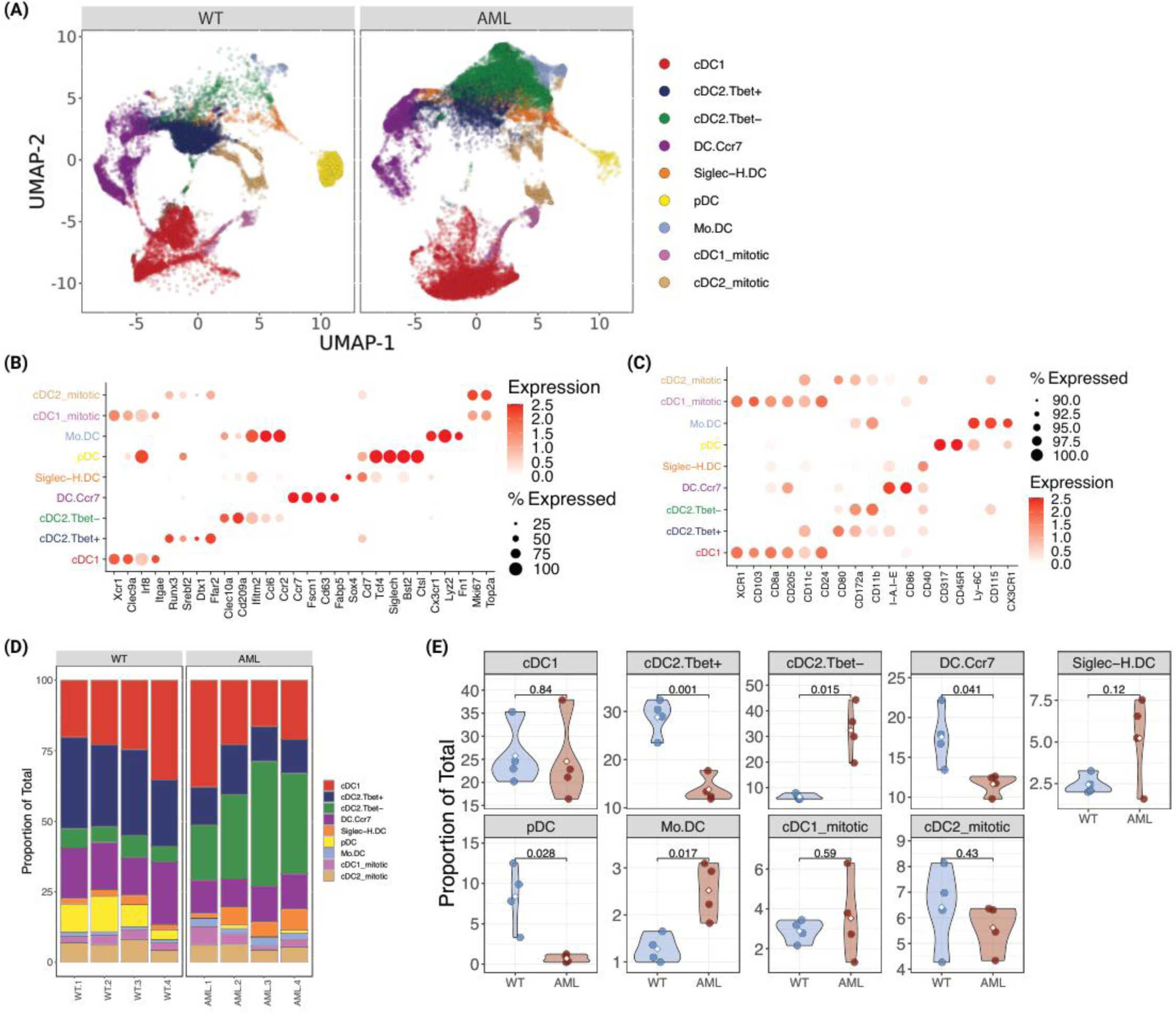
Single cell RNA-seq analysis of AML and WT mouse spleens identify changes in DC populations A. UMAP plot of DC subsets identified from WT and AML mice after magnetic bead enrichment for DCs. B. Mean expression of various DC gene markers across annotated DC subtypes. C. Mean abundance of protein markers across annotated DC subtypes. D. DC subtype proportions across samples AML (n=4) and WT (n=4). E. Violin plots of DC subtype proportions compared between WT and AML groups. Differences in means were determined using Student’s t-test.

### AML mice have disrupted CD4+ T cell phenotype and cytokines

It has been shown that mature DCs tune adaptive T cell responses into T-helper subsets (e.g. Th1, Th2, Th17, and Treg) [45–47] and given that the AML mice contain significantly increased levels of cDCs *in vivo* we hypothesized that CD4+ T cell populations would be altered systemically in the context of AML. We sampled peripheral blood from AML and WT healthy mice to measure the frequency of CD4+ T cells (Figure 4A). The frequency of circulating CD4+ T cells was not statistically different (Supplemental Figure 2) but when we compared the canonical T-helper transcription factors T-bet, GATA3, FOXP3, and RORγt we found that in AML mice they had increased Treg and Th17 populations (Figure 4B). Examination of the T-cell compartment identified from single-cell profiling of WT and AML splenocytes supported a strong shift away from naïve CD4+ T cells and an expansion of Tregs in AML spleen. Th17 cells could not be identified given the relatively low number of total T cells available for analysis (Supplemental Figure 7). The increased abundance of cDCs and the altered T-helper compartment led us to hypothesize that the cytokine profile in AML mice would also be altered. Serum samples from AML and WT healthy mice were analyzed by LegendPlex Mouse Inflammation Panel for inflammation associated cytokines (Figure 4C). We found that AML mice had increased levels of IL-1β and TNF-α, consistent with a chronic disease phenotype. We also observed increased levels of IL-27 which is a potent DC-secreted cytokine that can influence T cell responses after activation [48, 49] as well as the Treg and Th17 associated cytokines IL-10 and IL-17α (Figure 4C). Based on this finding we tested human FLT3-ITD+ AML patient samples by Luminex and saw significantly increased serum levels of IL-10 (Figure 4D) and IL-17A (Figure 4E) giving confidence that our mouse model is reflecting what is seen in AML patients. Together, these data suggest that in the context of AML, naïve CD4+ T cells are directed towards Th17 and Treg phenotypes by cDCs.

**Figure 4.**
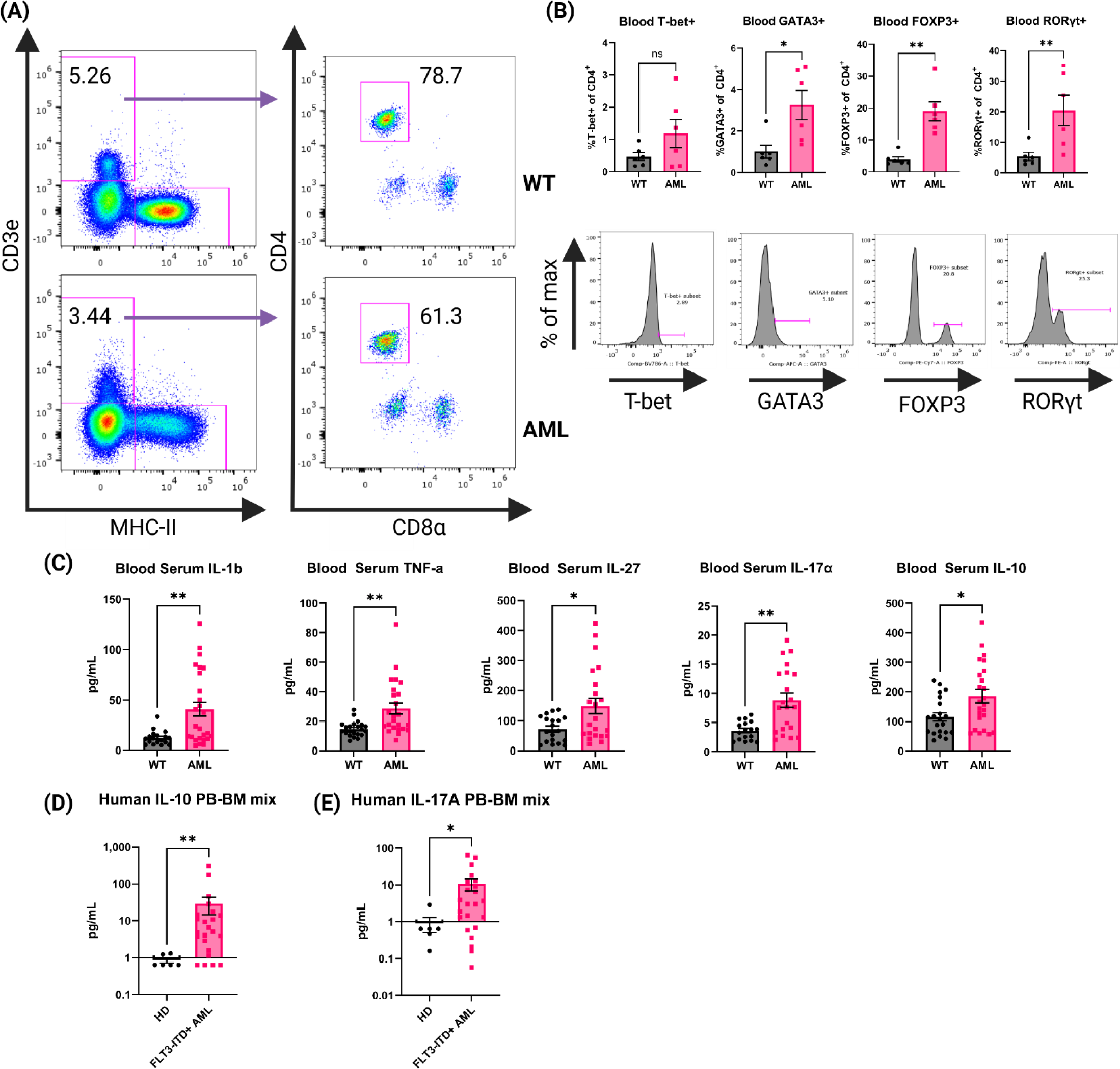
AML mouse blood exhibits a Th17 inflammatory phenotype A. Representative dot plots for gating on mouse blood CD4+ T cells by flow cytometry. Top row is WT and bottom row is AML. B. Transcription factor expression of CD4+ T cells. Top row is summary bar charts of transcription factor expression in CD4+ T cells from (A). Each symbol is an individual mouse. WT (n=6) and AML (n=6). Bottom row is representative histograms of each transcription factor. C. Blood serum cytokines in WT (n=21) and AML (24) mice. Each symbol is an individual mouse. Summary bar charts for selected inflammatory cytokines measured using BioLegend LegendPlex multiplex assay Mouse Inflammation Panel. D. Summary bar charts for Luminex measurement of human IL-10 cytokine detected in HD and AML patient samples of mixed peripheral blood (PB) and bone marrow (BM). HD (n=6) and FLT3-ITD+ AML (n=23). E. Summary bar charts for Luminex measurement of human IL-17 cytokine detected in HD and AML patient samples of mixed peripheral blood (PB) and bone marrow (BM). HD (n=6) and FLT3-ITD+ AML (n=23)

### AML mice support expansion of OT-II cells *in vivo* and AML DCs promote Th17 skewing *in vitro*

Given that we found the increase in T-bet- cDC2s and Th17 cells in AML mice, there was reason to investigate whether the CD4+ T cell phenotype was the result of interactions with DCs. We performed adoptive cell transfer (ACT) using naïve transgenic OT-II cells that are specific to OVA peptides [28, 50, 51] to study antigen specific activation of CD4+ T cells. Age-matched healthy WT or AML mice received equal numbers of naïve OT-II cells and one day following received an equal dose of whole-OVA protein. Spleens were harvested for flow cytometric analysis 10 days after OVA injection (Figure 5A). Transferred cells were identified using congenic markers (Figure 5B). When probed for transcription factors FOXP3 and RORγt we did not see expression of either transcription factor. We did observe significantly more CD44+ OT-II cells and with increased frequency in AML host mice at Day 11 suggesting that AML are more supportive of CD4+ T cells after transfer (Figure 5C). Since we did not see evidence of specific Th skewing *in vivo* we next tested an *in vitro* approach where isolated cDCs and naïve OT-II T cells were co-cultured with OVA323 peptide (Figure 5D). After five days of culture we analyzed secretion of IL-17A by flow cytometry (Figure 5E). In the groups that had the addition of the OT-II dominant antigen OVA323, we saw an increased trend of IL-17A secretion in the AML cDC co-cultures (Figure 5F). Taken together our data suggests that AML mice have a DC phenotype that supports CD4+ T cell retention and polarization of naïve CD4+ T cells into a Th17 phenotype. In summary, we find that in the context of our AML mouse model, DCs inherit the FLT3-ITD mutation and DC homeostasis is disrupted. We also report that CD4+ T cells are skewed towards Treg and Th17 phenotypes *in vivo* and adoptively transferred OT-II cells are retained in higher numbers in AML recipients. *In vitro* co-cultures of OT-II cells and DCs resulted in more IL-17A secretion in the AML samples. All of these data together suggest that the increased DC phenotype observed also directly impacts the CD4+ T cell compartment, which results in a microenvironment that is tumor supportive.

**Figure 5.**
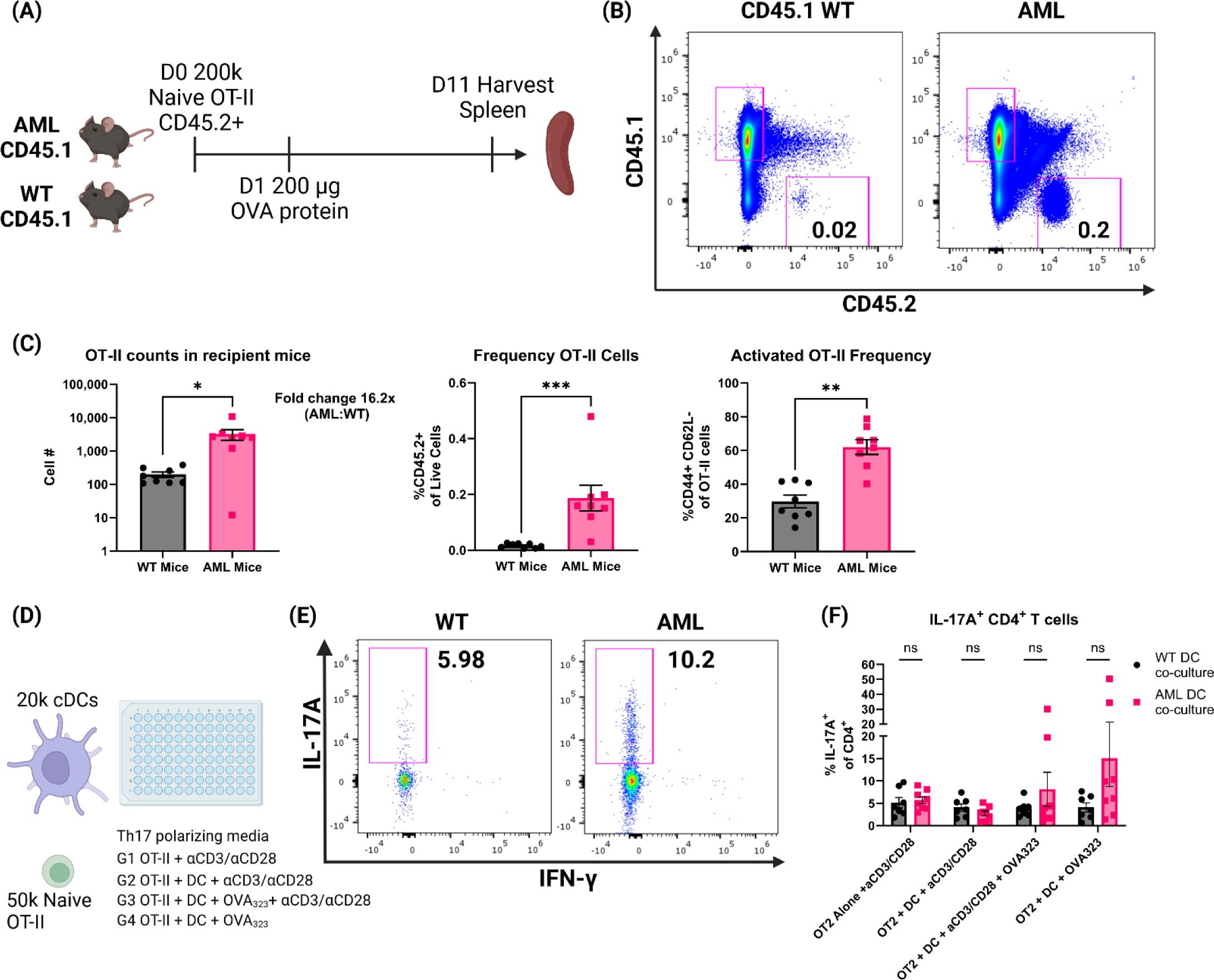
AML mice retain more OT-II cells and AML DCs promote IL-17A production A. Experimental outline of OT-II adoptive cell transfer. CD45.1+ AML (n=8) and WT (n=8) mice are injected intravenously with 200k naïve CD45.2+ OT-II cells on Day 0. 24 hours later mice were injected intraperitonealy with 200 µg whole OVA protein. 10 days later mice were sacrificed and spleens were harvested for flow cytometry analysis of remaining CD45.2+ OT-II cells. B. Representative dot plots of spleen resident CD45.2+ OT-II cells by flow cytometry. Frequency of live single cells noted in the plots. C. Summary bar charts of OT-II cell counts (left panel) and frequency of live single cells (right panel). Each symbol is an individual mouse and summary data collected from two experiments. AML (n=8) and WT (n=8). D. Experimental outline of *in vitro* co-cultures of OT-II cells and cDCs. Naïve OT-II cells were magnetically sorted from spleen tissue using BioLegend MojoSort Mouse CD4 Naïve T Cell Isolation Kit. After magnetic sorting samples were verified for purity by flow cytometry. cDCs were magnetically isolated using BioLegend MojoSort Mouse CD11c+ beads. After magnetic sorting samples were verified by flow cytometry. OT-II cells were plated at 50k cells per well and cDCs were plated at 20k cells per well for five days in Th17 polarizing media using CellXVivo Mouse Th17 Cell Differentiation Kit CDK017 from R&D Systems. E. Representative dot plots of IL-17A+ OT-II cells by flow cytometry. Frequency of CD4+ cells noted in the plots. F. Summary bar charts of IL-17A+ CD4+ T cells from (E). Each symbol is an individual sample and summary data collected from four experiments. AML (n=8) and WT (n=7).

## Discussion

FLT3-ITD is one of the most common mutations found in AML patients and research focus has been placed mainly on the AML blasts. DCs are the most responsive cell type to FLT3 signaling and they rely on this pathway for development [52, 53]. We found that in bone marrow from FLT3-ITD+ AML patients, the frequency of cDCs was heterogeneous and patients could be stratified into high/medium/low frequencies. FLT3-ITD- bone marrow samples also had elevated levels of cDCs in the bone marrow compared to HD, but the highest observed frequencies were in the FLT3-ITD+ AML, suggesting that there are cDC precursors that are not tumor blasts that retain this mutation and expand. Our data shows that in AML the cDC1 and cDC2 subsets are disrupted and the frequency of XCR1/cDC1 double-negative cDCs was significantly higher in FLT3-ITD+ samples. This may suggest that patients with FLT3-ITD+ AML may retain more precursors than fully committed DC cells. Reports on bone marrow frequency of cDCs in the literature are sparse [54] but what we observe is consistent with the data reported, where a subset of AML patients have increased levels of cDCs in their bone marrow. However, that study did not have FLT3-ITD mutation status for patients. The majority of AML-DC reports focus on measuring circulating pDCs and cDCs [23, 55, 56]support the idea that FLT3-ITD+ AML patients have increased DC frequency in the peripheral blood. Immunophenotyping more FLT3-ITD+ AML patients will be important for understanding the changes to DC subsets in the bone marrow and the periphery.

We found that our spontaneous AML mice had a significant DC phenotype. It has been reported previously in a non-AML context that the FLT3-ITD mutation led to cDC expansion systemically [25]. We see that in the context of AML the expansion of cDCs is even more profound and was a result of FLT3-ITD *in vivo*. This suggests that in our model, the FLT3-ITD mutation drives a proportion of DC-precursors into mature DCs rather than malignant cells. The molecular understanding of DCs has increased significantly due to advances in sequencing technologies [57–59]. We used the models from Brown et. al. to inform our scRNA-seq analysis [5]. To the best of our knowledge, our report is the first to provide scRNA-seq data on DCs in the context of a model of FLT3-ITD+ AML. In combination with our ADT data, we can confidently identify *bona fide* DCs in the spleen. This has allowed us to more accurately describe changes to the DC population without contamination from other myeloid subsets and tumor blasts that have a shared lineage and could confound our DC findings. Our AML mice had more T-bet^-^ cDC2s, a population of cells that have been shown to promote polarization of naïve CD4+ T cells into Th17 phenotypes [5]. This finding was unexpected to us because reports suggest that strong FLT3 signaling *in vivo* drives all DC subsets to expand [9, 25, 60]. This T-bet^-^ cDC2 cell type is novel and will be the subject of future studies to understand how it is functionally different from other DCs in AML. Our data is concordant with what was reported by Brown et. al., which could have clinical implications for AML patients. The role of Th17 cells in cancer is controversial but in AML they have been found to be deleterious for patient survival [43, 44, 61, 62]. Our mouse model of AML recapitulates findings from humans where the inflammatory cytokine milieu is permissive to Th17 polarization [63, 64]. [36, 37, 65, 66] We recognize that the DC phenotype we describe in our human samples is different from the mouse DC phenotype. This was not unexpected as AML is a very genetically complicated cancer [14, 15]. Our mouse model has a select number of mutations (FLT3-ITD, TET2 KO, and TP53KO) that leads to AML-like disease but is a significantly smaller list of mutations compared to human AML patients [14]. It is likely that the increased complexity of mutations in the AML patient samples is impacting DC subsets in a way that we cannot replicate in the GEMM that we use. What may address this discrepancy is the use of ATACseq and scRNA-seq on both mouse and human AML DCs to compare their chromatin accessibility and transcriptomes. Such data would shed light on what differences and parallels exists between species regarding the active sites of transcription and gene expression as a result of inherited leukemia mutations.

It is important in our view to consider the contribution of DCs on the T cell compartment as DCs are continuously supplied to the periphery and difficult to ablate compared to lymphocytes. In summary, we have shown that in some subsets of FLT3-ITD+ AML patients the frequency of bone marrow DCs is increased. Using a mouse model of AML we characterized this increase in DC abundance as a result of FLT3-ITD and that they are skewed towards a T-bet^-^ cDC2 phenotype. These cells are adept at polarizing naïve CD4+ T cells into the Th17 subset. This work provides novel scRNA-seq data on DCs and further supports rationale for the need to improve the characterization and function of non-tumor myeloid cells and their impact on anti-tumor responses. This is particularly important with the growing interest in developing immunotherapies for AML [36, 37, 65, 66].

## Supporting information

Supplemental Data PAF AML DC

## Acknowledgments

Flow cytometry was performed at the OHSU Flow Cytometry Core Facility. Sequencing assays were performed by the OHSU Massively Parallel Sequencing Shared Resource.

## Bibliography

1. Steinman, R.M. and Z.A. Cohn, Identification of a novel cell type in peripheral lymphoid organs of mice. I. Morphology, quantitation, tissue distribution. J Exp Med, 1973. 137(5): p. 1142–62.

2. Steinman, R.M., et al., Identification of a novel cell type in peripheral lymphoid organs of mice. V. Purification of spleen dendritic cells, new surface markers, and maintenance in vitro. J Exp Med, 1979. 149(1): p. 1–16.

3. Leylek, R., et al., Integrated Cross-Species Analysis Identifies a Conserved Transitional Dendritic Cell Population. Cell Rep, 2019. 29(11): p. 3736–3750 e8.

4. Ma, W., et al., Single cell RNA-Seq reveals pre-cDCs fate determined by transcription factor combinatorial dose. BMC Mol Cell Biol, 2019. 20(1): p. 20.

5. Brown, C.C., et al., Transcriptional Basis of Mouse and Human Dendritic Cell Heterogeneity. Cell, 2019. 179(4): p. 846–863 e24.

6. Cabeza-Cabrerizo, M., et al., Dendritic Cells Revisited. Annu Rev Immunol, 2021. 39: p. 131–166.

7. Murphy, T.L., et al., Transcriptional Control of Dendritic Cell Development. Annu Rev Immunol, 2016. 34: p. 93–119.

8. Anderson, D.A., 3rd, K.M. Murphy, and C.G. Briseno, *Development, Diversity, and Function of Dendritic Cells in Mouse and Human*. Cold Spring Harb Perspect Biol, 2018. 10(11).

9. Laouar, Y., et al., STAT3 is required for Flt3L-dependent dendritic cell differentiation. Immunity, 2003. 19(6): p. 903–12.

10. Maraskovsky, E., et al., Dramatic increase in the numbers of functionally mature dendritic cells in Flt3 ligand-treated mice: multiple dendritic cell subpopulations identified. J Exp Med, 1996. 184(5): p. 1953–62.

11. Pulendran, B., et al., Developmental pathways of dendritic cells in vivo: distinct function, phenotype, and localization of dendritic cell subsets in FLT3 ligand-treated mice. J Immunol, 1997. 159(5): p. 2222–31.

12. Shurin, M.R., et al., FLT3 ligand induces the generation of functionally active dendritic cells in mice. Cell Immunol, 1997. 179(2): p. 174–84.

13. Ferrara, F. and C.A. Schiffer, Acute myeloid leukaemia in adults. Lancet, 2013. 381(9865): p. 484–95.

14. Tyner, J.W., et al., Functional genomic landscape of acute myeloid leukaemia. Nature, 2018. 562(7728): p. 526–531.

15. Estey, E. and H. Dohner, Acute myeloid leukaemia. Lancet, 2006. 368(9550): p. 1894–907.

16. Patel, J.P., et al., Prognostic relevance of integrated genetic profiling in acute myeloid leukemia. N Engl J Med, 2012. 366(12): p. 1079–89.

17. Nakao, M., et al., Internal tandem duplication of the flt3 gene found in acute myeloid leukemia. Leukemia, 1996. 10(12): p. 1911–8.

18. Kiyoi, H., et al., Internal tandem duplication of the FLT3 gene is a novel modality of elongation mutation which causes constitutive activation of the product. Leukemia, 1998. 12(9): p. 1333–7.

19. Yokota, S., et al., Internal tandem duplication of the FLT3 gene is preferentially seen in acute myeloid leukemia and myelodysplastic syndrome among various hematological malignancies. A study on a large series of patients and cell lines. Leukemia, 1997. 11(10): p. 1605–9.

20. Gilliland, D.G. and J.D. Griffin, The roles of FLT3 in hematopoiesis and leukemia. Blood, 2002. 100(5): p. 1532–42.

21. Lee, B.H., et al., FLT3 mutations confer enhanced proliferation and survival properties to multipotent progenitors in a murine model of chronic myelomonocytic leukemia. Cancer Cell, 2007. 12(4): p. 367–80.

22. Mohty, M., et al., Circulating blood dendritic cells from myeloid leukemia patients display quantitative and cytogenetic abnormalities as well as functional impairment. Blood, 2001. 98(13): p. 3750–6.

23. Ma, L., et al., Circulating myeloid and lymphoid precursor dendritic cells are clonally involved in myelodysplastic syndromes. Leukemia, 2004. 18(9): p. 1451–6.

24. Kurtz, K.J., et al., Murine Models of Acute Myeloid Leukemia. Front Oncol, 2022. 12: p. 854973.

25. Lau, C.M., et al., Leukemia-associated activating mutation of Flt3 expands dendritic cells and alters T cell responses. J Exp Med, 2016. 213(3): p. 415–31.

26. Kelly, L.M., et al., FLT3 internal tandem duplication mutations associated with human acute myeloid leukemias induce myeloproliferative disease in a murine bone marrow transplant model. Blood, 2002. 99(1): p. 310–8.

27. Moran-Crusio, K., et al., Tet2 loss leads to increased hematopoietic stem cell self-renewal and myeloid transformation. Cancer Cell, 2011. 20(1): p. 11–24.

28. Barnden, M.J., et al., Defective TCR expression in transgenic mice constructed using cDNA-based alpha- and beta-chain genes under the control of heterologous regulatory elements. Immunol Cell Biol, 1998. 76(1): p. 34–40.

29. Butler, A., et al., Integrating single-cell transcriptomic data across different conditions, technologies, and species. Nat Biotechnol, 2018. 36(5): p. 411–420.

30. Wolock, S.L., R. Lopez, and A.M. Klein, Scrublet: Computational Identification of Cell Doublets in Single-Cell Transcriptomic Data. Cell Syst, 2019. 8(4): p. 281–291 e9.

31. Young, M.D. and S. Behjati, SoupX removes ambient RNA contamination from droplet-based single-cell RNA sequencing data. Gigascience, 2020. 9(12).

32. Finak, G., et al., MAST: a flexible statistical framework for assessing transcriptional changes and characterizing heterogeneity in single-cell RNA sequencing data. Genome Biol, 2015. 16: p. 278.

33. Aran, D., et al., Reference-based analysis of lung single-cell sequencing reveals a transitional profibrotic macrophage. Nat Immunol, 2019. 20(2): p. 163–172.

34. Andreatta, M., et al., Interpretation of T cell states from single-cell transcriptomics data using reference atlases. Nat Commun, 2021. 12(1): p. 2965.

35. Bottomly, D., et al., Integrative analysis of drug response and clinical outcome in acute myeloid leukemia. Cancer Cell, 2022. 40(8): p. 850–864 e9.

36. Moshofsky, K.B., et al., Acute myeloid leukemia-induced T-cell suppression can be reversed by inhibition of the MAPK pathway. Blood Adv, 2019. 3(20): p. 3038–3051.

37. Romine, K.A., et al., BET inhibitors rescue anti-PD1 resistance by enhancing TCF7 accessibility in leukemia-derived terminally exhausted CD8(+) T cells. Leukemia, 2023. 37(3): p. 580–592.

38. Marino, S., et al., Induction of medulloblastomas in p53-null mutant mice by somatic inactivation of Rb in the external granular layer cells of the cerebellum. Genes Dev, 2000. 14(8): p. 994–1004.

39. Clausen, B.E., et al., Conditional gene targeting in macrophages and granulocytes using LysMcre mice. Transgenic Res, 1999. 8(4): p. 265–77.

40. Shih, A.H., et al., Mutational cooperativity linked to combinatorial epigenetic gain of function in acute myeloid leukemia. Cancer Cell, 2015. 27(4): p. 502–15.

41. Rosnet, O., et al., Human FLT3/FLK2 receptor tyrosine kinase is expressed at the surface of normal and malignant hematopoietic cells. Leukemia, 1996. 10(2): p. 238–48.

42. Drexler, H.G., Expression of FLT3 receptor and response to FLT3 ligand by leukemic cells. Leukemia, 1996. 10(4): p. 588–99.

43. Han, Y., et al., Th17 cells and interleukin-17 increase with poor prognosis in patients with acute myeloid leukemia. Cancer Sci, 2014. 105(8): p. 933–42.

44. Bailey, S.R., et al., Th17 cells in cancer: the ultimate identity crisis. Front Immunol, 2014. 5: p. 276.

45. Oth, T., et al., Monitoring the initiation and kinetics of human dendritic cell-induced polarization of autologous naive CD4+ T cells. PLoS One, 2014. 9(8): p. e103725.

46. Peters, M., et al., T-cell polarization depends on concentration of the danger signal used to activate dendritic cells. Immunol Cell Biol, 2010. 88(5): p. 537–44.

47. Boonstra, A., et al., Flexibility of mouse classical and plasmacytoid-derived dendritic cells in directing T helper type 1 and 2 cell development: dependency on antigen dose and differential toll-like receptor ligation. J Exp Med, 2003. 197(1): p. 101–9.

48. Ahmadi, F., et al., cDC1-derived IL-27 regulates small intestinal CD4+ T cell homeostasis in mice. J Exp Med, 2023. 220(3).

49. Pennock, N.D., L. Gapin, and R.M. Kedl, IL-27 is required for shaping the magnitude, affinity distribution, and memory of T cells responding to subunit immunization. Proc Natl Acad Sci U S A, 2014. 111(46): p. 16472–7.

50. Robertson, J.M., P.E. Jensen, and B.D. Evavold, DO11.10 and OT-II T cells recognize a C-terminal ovalbumin 323- 339 epitope. J Immunol, 2000. 164(9): p. 4706–12.

51. Yang, L., et al., Generation of functional antigen-specific T cells in defined genetic backgrounds by retrovirus-mediated expression of TCR cDNAs in hematopoietic precursor cells. Proc Natl Acad Sci U S A, 2002. 99(9): p. 6204–9.

52. Waskow, C., et al., The receptor tyrosine kinase Flt3 is required for dendritic cell development in peripheral lymphoid tissues. Nat Immunol, 2008. 9(6): p. 676–83.

53. Durai, V., et al., Altered compensatory cytokine signaling underlies the discrepancy between Flt3(-/-) and Flt3l(-/-) mice. J Exp Med, 2018. 215(5): p. 1417–1435.

54. Derolf, A.R., et al., Dendritic cells in bone marrow at diagnosis and after chemotherapy in adult patients with acute myeloid leukaemia. Scand J Immunol, 2014. 80(6): p. 424–31.

55. Rickmann, M., et al., Elevated frequencies of leukemic myeloid and plasmacytoid dendritic cells in acute myeloid leukemia with the FLT3 internal tandem duplication. Ann Hematol, 2011. 90(9): p. 1047–58.

56. Rickmann, M., et al., Monitoring dendritic cell and cytokine biomarkers during remission prior to relapse in patients with FLT3-ITD acute myeloid leukemia. Ann Hematol, 2013. 92(8): p. 1079–90.

57. Sichien, D., et al., IRF8 Transcription Factor Controls Survival and Function of Terminally Differentiated Conventional and Plasmacytoid Dendritic Cells, Respectively. Immunity, 2016. 45(3): p. 626–640.

58. Balan, S., et al., Large-Scale Human Dendritic Cell Differentiation Revealing Notch-Dependent Lineage Bifurcation and Heterogeneity. Cell Rep, 2018. 24(7): p. 1902–1915 e6.

59. Kirkling, M.E., et al., Notch Signaling Facilitates In Vitro Generation of Cross-Presenting Classical Dendritic Cells. Cell Rep, 2018. 23(12): p. 3658–3672 e6.

60. Lin, D.S., et al., Single-cell analyses reveal the clonal and molecular aetiology of Flt3L-induced emergency dendritic cell development. Nat Cell Biol, 2021. 23(3): p. 219–231.

61. Barilla, R.M., et al., Specialized dendritic cells induce tumor-promoting IL-10(+)IL-17(+) FoxP3(neg) regulatory CD4(+) T cells in pancreatic carcinoma. Nat Commun, 2019. 10(1): p. 1424.

62. Musuraca, G., et al., IL-17/IL-10 double-producing T cells: new link between infections, immunosuppression and acute myeloid leukemia. J Transl Med, 2015. 13: p. 229.

63. Sanchez-Correa, B., et al., Cytokine profiles in acute myeloid leukemia patients at diagnosis: survival is inversely correlated with IL-6 and directly correlated with IL-10 levels. Cytokine, 2013. 61(3): p. 885–91.

64. Carey, A., et al., Identification of Interleukin-1 by Functional Screening as a Key Mediator of Cellular Expansion and Disease Progression in Acute Myeloid Leukemia. Cell Rep, 2017. 18(13): p. 3204–3218.

65. Lamble, A.J., et al., Reversible suppression of T cell function in the bone marrow microenvironment of acute myeloid leukemia. Proc Natl Acad Sci U S A, 2020. 117(25): p. 14331–14341.

66. Lamble, A.J. and E.F. Lind, Targeting the Immune Microenvironment in Acute Myeloid Leukemia: A Focus on T Cell Immunity. Front Oncol, 2018. 8: p. 213.

