## Supplemental Data PAF AML DC for "Leukemic mutation FLT3-ITD is retained in dendritic cells and disrupts their homeostasis leading to expanded Th17 frequency"

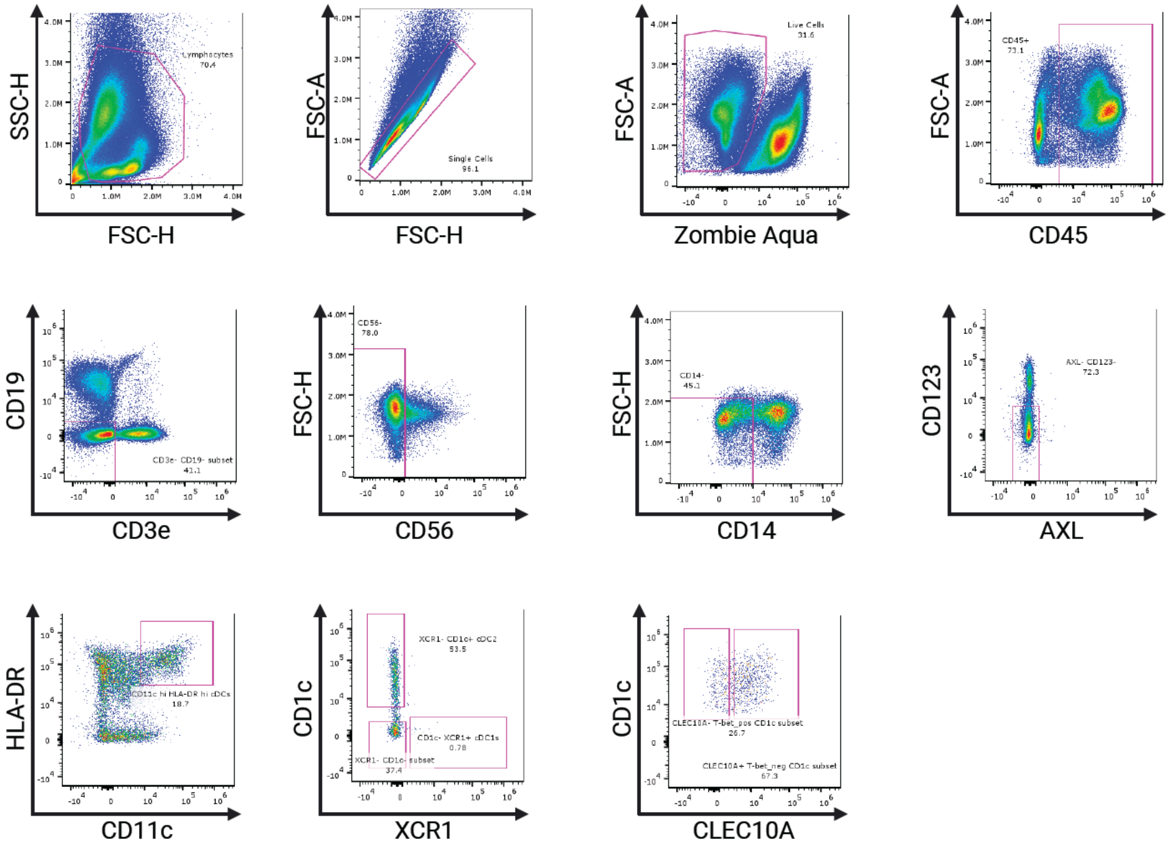


Supplemental Figure 1. Gating scheme for human bone marrow flow cytometry and DC-like score

A. FlowJo gating scheme showing identification of human cDCs from bone marrow aspirate.
B. Dot plots showing DC Cell-Type scores from AML patient samples that were identified using Weighted Gene Co-expression Network Analysis (WGCNA) from BEAT AML Bottomly et. al. 2022 https://doi.org/10.1016/j.ccell.2022.07.002.

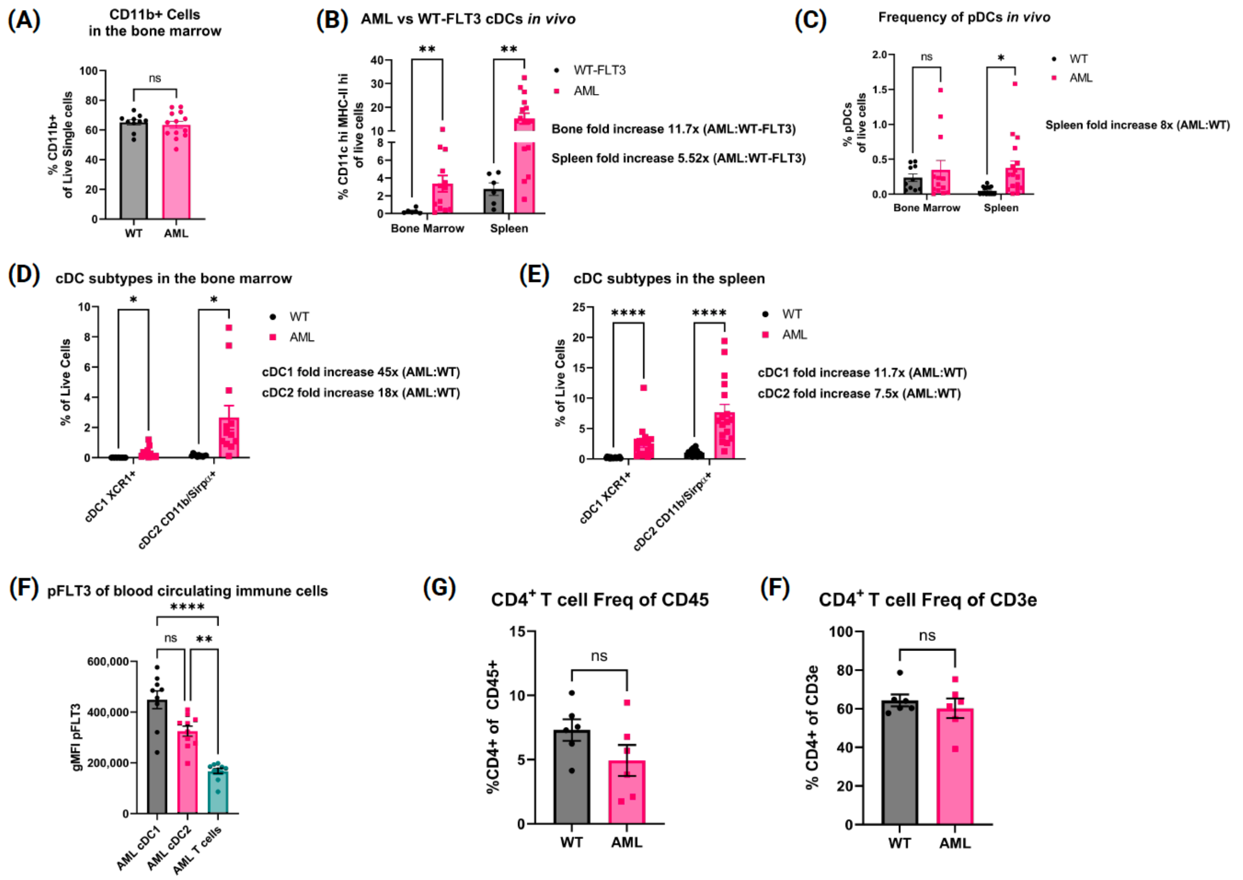


Supplemental Figure 2. Mouse Model of AML Has Significantly Increased cDCs In Vivo

A. Summary bar chart of CD11b+ cell frequency in the bone marrow of WT (n=10) and AML (n=13) mice.
B. Summary bar chart of cDC frequency in the bone marrow and spleen compartments of WT-FLT3 LysM-Cre+ TET2loxp/loxp mice (n=6) and AML mice (n= bone marrow n=13 AML spleens n=16).
C. Summary bar chart of pDC frequency in bone marrow and spleen. Each symbol is one mouse. WT bone marrow n=10 WT spleens n=23. AML bone marrow n=13 AML spleens n=16.
D. Summary bar chart of cDC1 and cDC2 frequency in bone marrow. Each symbol is one mouse. WT bone marrow n=10. AML bone marrow n=13.
E. Summary bar chart of cDC1 and cDC2 frequency in spleens. Each symbol is one mouse. WT bone marrow n=23. AML bone marrow n=16
F. Summary bar chart of blood circulating AML cDC1 and AML cDC2 and AML T cells gMFI pFLT3. N=10.

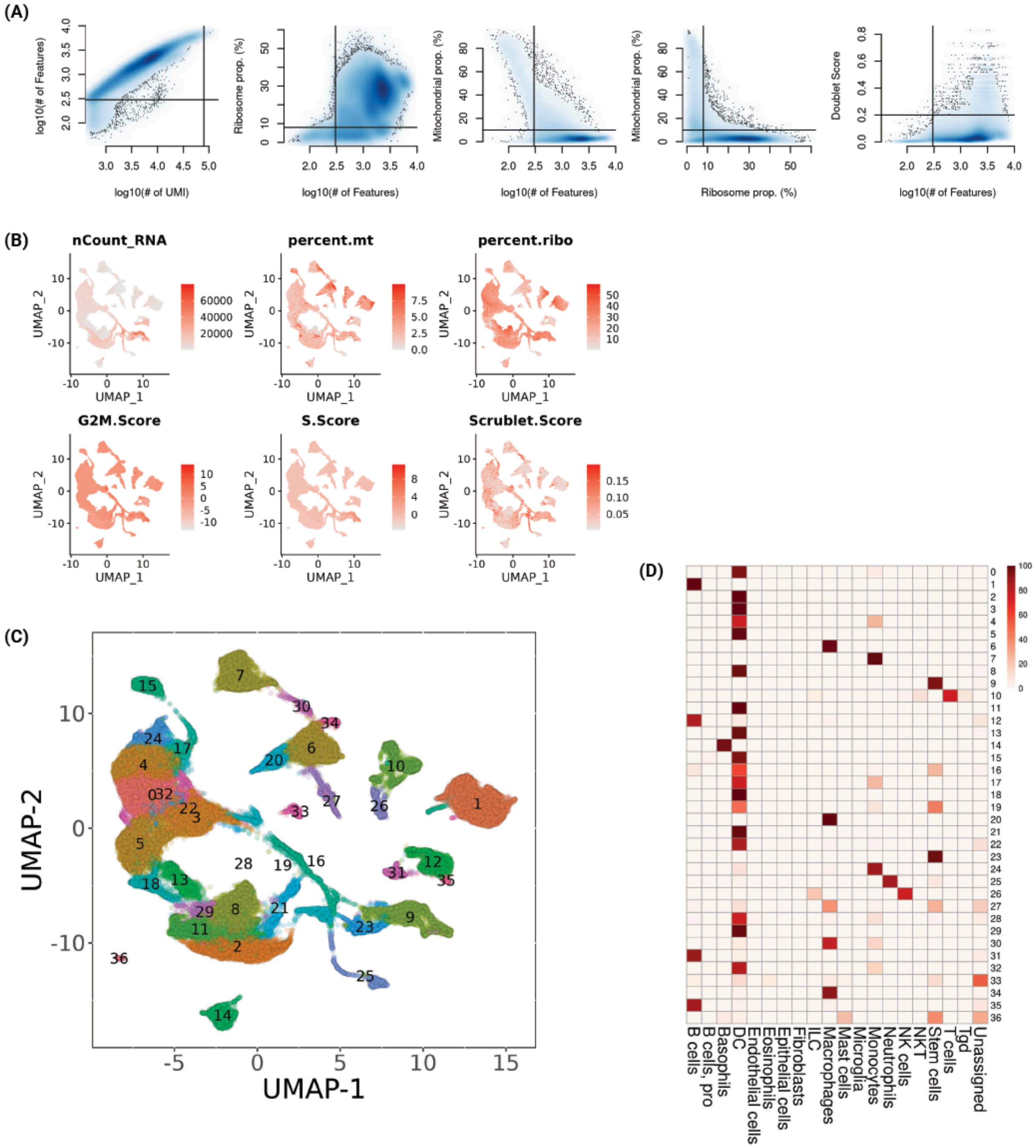


Supplemental Figure 3. Mouse scRNA-seq QC and processing metrics

A. Filtering criteria used to isolate high-quality cells for downstream analysis, including number of detectible features/UMI, ribosomal and mitochondrial gene proportions, and Doublet Score.
B. QC metrics and cell-cycle scoring shown for all cells.
C. UMAP representation of all high-quality cells. Colors represent final clusters determined by unsupervized clustering analysis.
D. Proportion of each cluster annotating to major lineages determined by SingleR supervised classification against the Immgen Database.

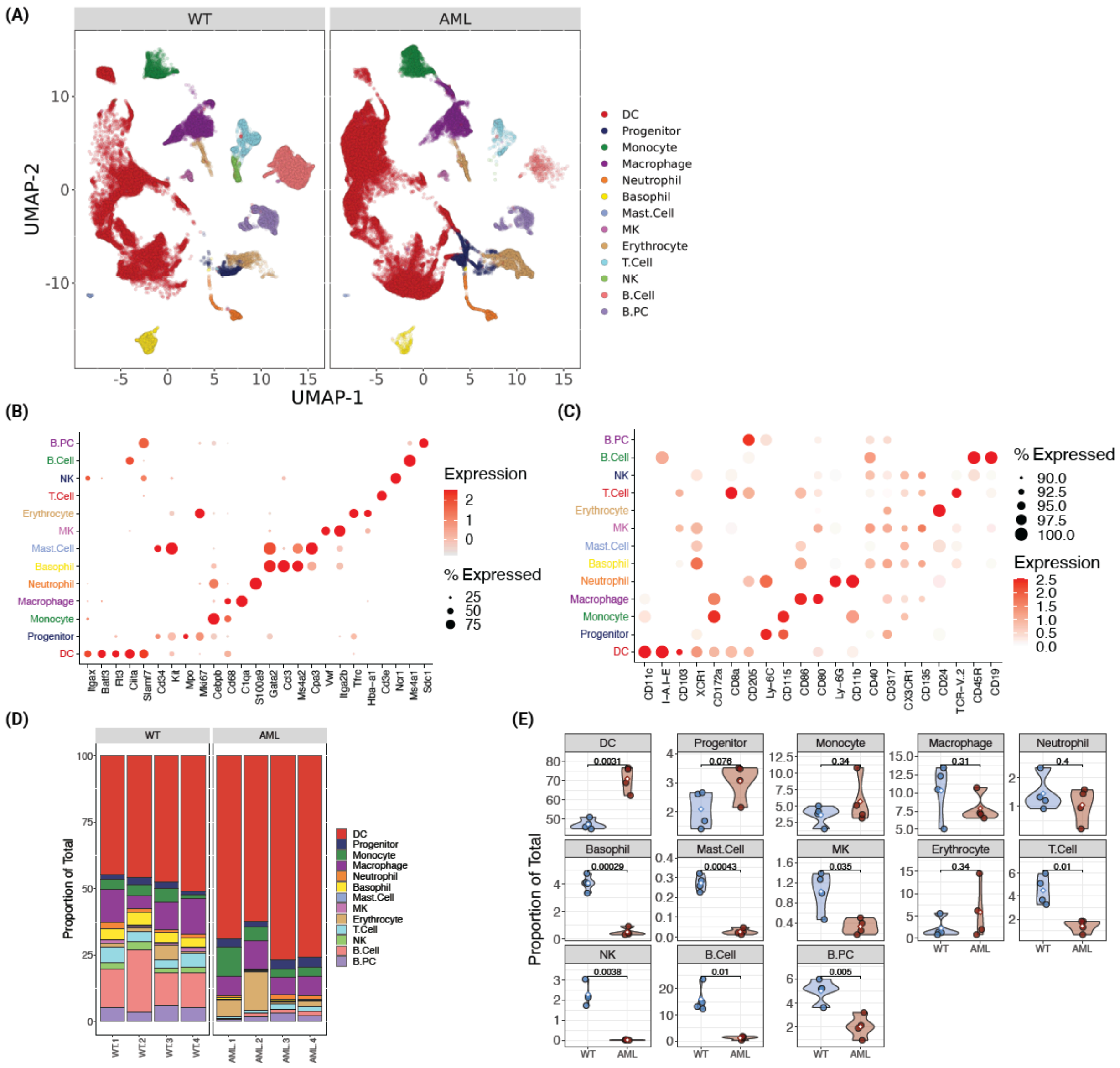

Supplemental Figure 4: Single cell RNA-seq profiling of AML and WT mouse spleens.

A. UMAP plot of total cells derived from WT and AML mice after magnetic bead enrichment for DCs.
B. Mean expression of various linage markers across annotated cell types.
C. Mean abundance of protein markers across annotated cell types.
D. Cell type proportions across samples AML (n=4) and WT (n=4).
E. Violin plots of cell type proportions compared between WT and AML groups. Differences in means were determined using Student's t-test.

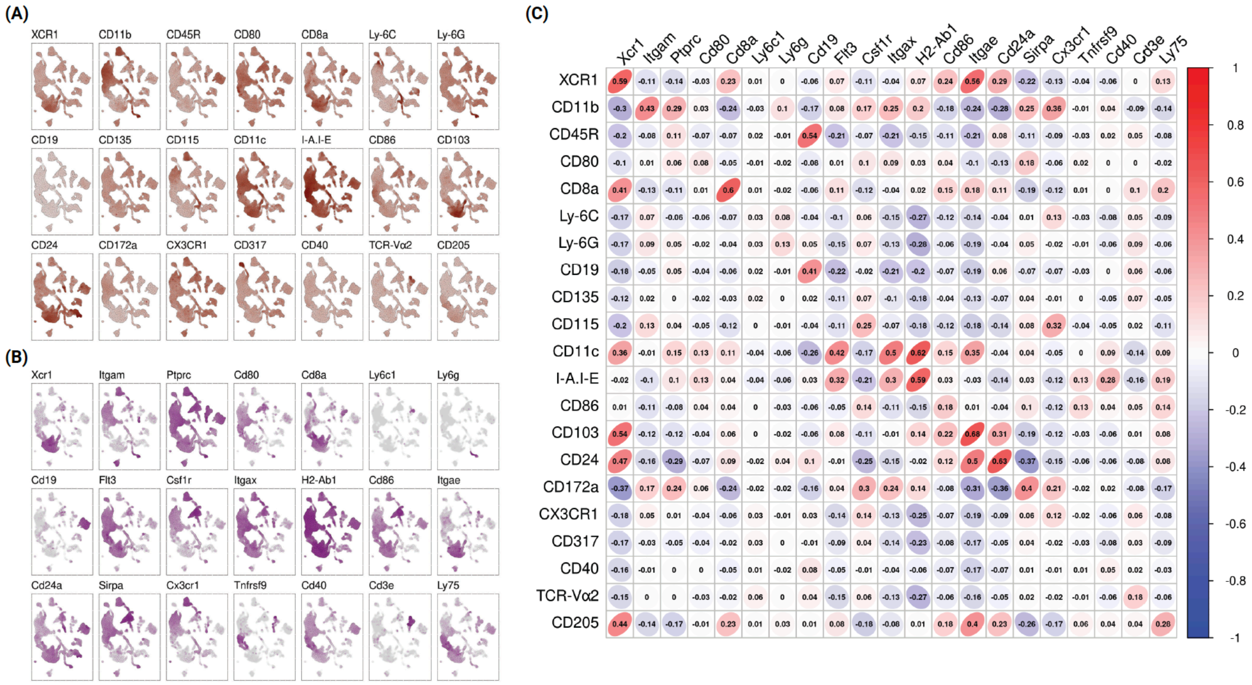


Supplemental Figure 5. Comparison of GEX and Protein ADT profiles across single-cells

A. UMAP representation of for protein ADT abundances.
B. UMAP representation of gene expression markers associated with proteins profiled in A.
C. Correlation matrix for surface proteins and corresponding transcripts across all cells.

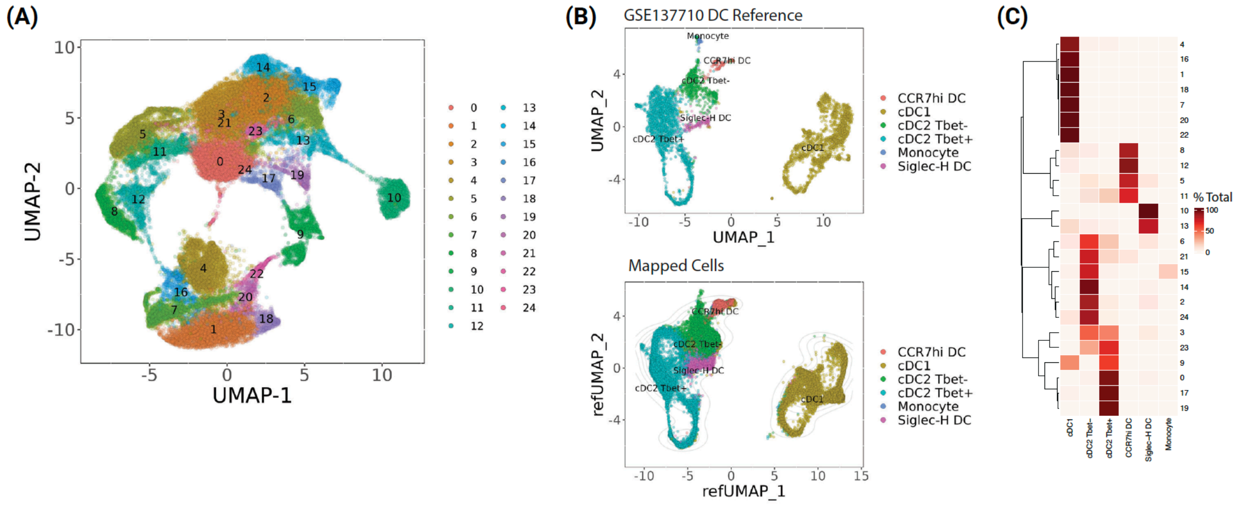


Supplemental Figure 6. Reference based mapping delineates DC heterogeneity in WT and AML mouse spleens.

A. UMAP visualization of the identified DC compartment. Colors represent final clusters determined by unsupervized clustering analysis.
B. Upper panel, GSE137710 DC Reference UMAP from re-analysis of data derived from Brown *et. al.* (GSE137710). Bottom panel, AML and WT mouse splenocytes mapped to Brown *et. al.* reference.
C. Proportion of cells within each cluster annotating to various DC subtypes after classification from reference based mapping.

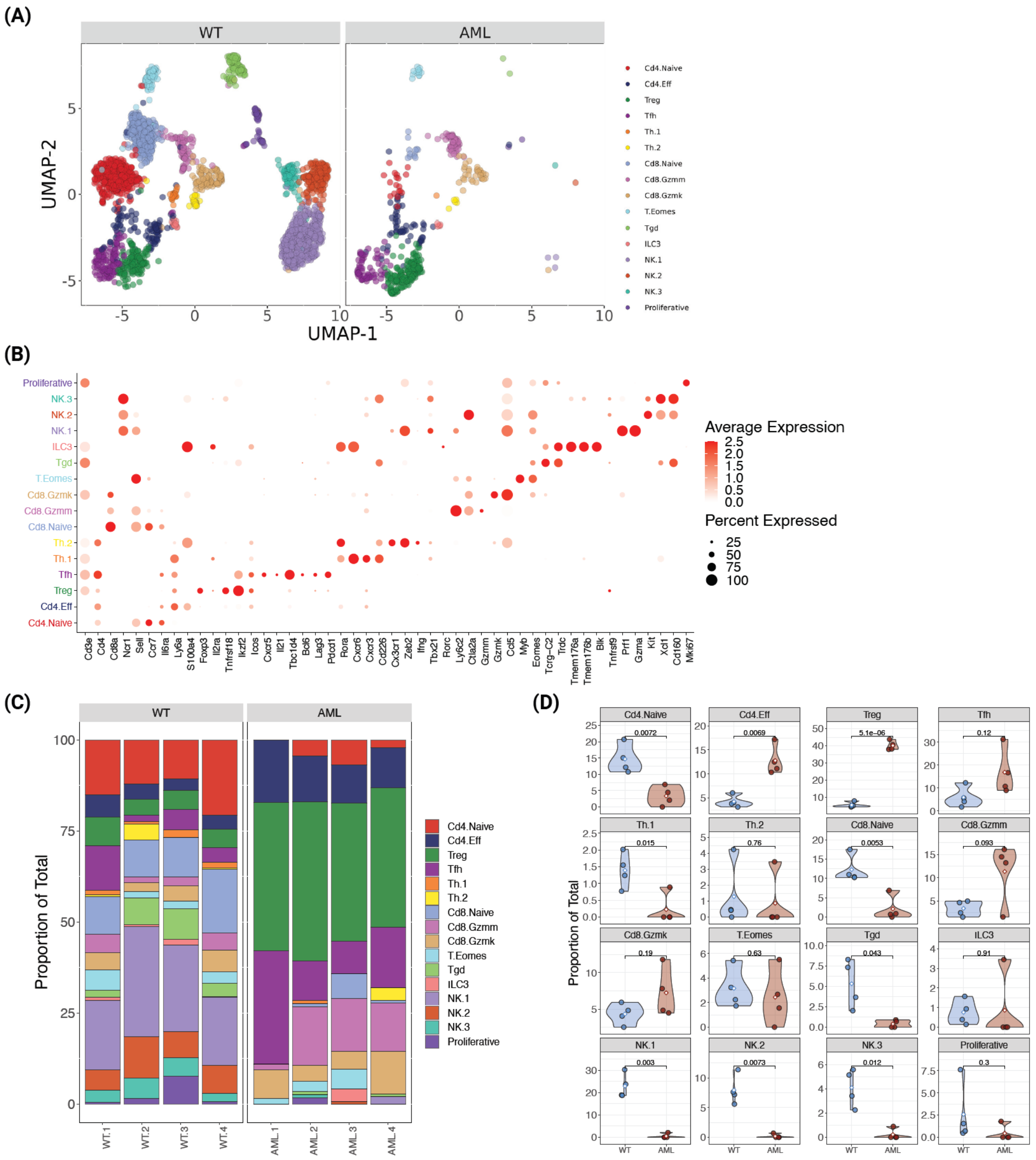

Supplemental Figure 7: Single cell RNA-seq analysis of AML and WT mouse spleens identify changes in T-cell phenotypes.

A. UMAP plot of T-cell and NK subsets identified from WT and AML mice.
B. Mean expression of various T and NK cell gene markers across annotated T-cell subtypes.
C. T and NK cell subtype proportions across samples AML (n=4) and WT (n=4).
D. Violin plots of T and NK cell subtype proportions compared between WT and AML groups. Differences in means were determined using Student's t-test.
